# ICELLNET: a transcriptome-based framework to dissect intercellular communication

**DOI:** 10.1101/2020.03.05.976878

**Authors:** Floriane Noël, Lucile Massenet-Regad, Irit Carmi-Levy, Antonio Cappuccio, Maximilien Grandclaudon, Coline Trichot, Yann Kieffer, Fatima Mechta-Grigoriou, Vassili Soumelis

## Abstract

Cell-to-cell communication can be inferred from ligand-receptor expression in cell transcriptomic datasets. However, important challenges remain: 1) global integration of cell-to-cell communication, 2) biological interpretation, and 3) application to individual cell population transcriptomic profiles. We developed ICELLNET, a transcriptomic-based framework integrating: 1) an original expert-curated database of ligand-receptor interactions accounting for multiple subunits expression, 2) quantification of communication scores, 3) the possibility to connect a cell population of interest with 31 reference human cell types (BioGPS), and 4) three visualization modes to facilitate biological interpretation. We applied ICELLNET to uncover different communication in breast cancer associated fibroblast (CAF) subsets. ICELLNET also revealed autocrine IL-10 as a switch to control human dendritic cell communication with up to 12 other cell types, four of which were experimentally validated. In summary, ICELLNET is a global, versatile, biologically validated, and easy-to-use framework to dissect cell communication from single or multiple cell-based transcriptomic profile(s).

## Introduction

Cell-to-cell communication is at the basis of the higher order organization observed in tissues, organs, and organisms, at steady-state and in response to stress. It involves a “messenger” or “sender” cell, which transmits information signals to a “receiving” or “target” cell. Information is generally coded in the form of a chemical molecule that is sensed by the target cell through a cognate receptor. Multiple cells or cell types communicating with each other form cell communication networks.

In mammalian organisms, endocrine communication involves cells that may be at very distant anatomical sites. However, cell communication also takes place locally through cell-to-cell contacts, or through inflammatory molecules. Cytokines and other mediators can be involved in distant as well as local communication^1–3^. Hence, when deciphering cell-to-cell communication, one should account for potential signals coming both from spatially proximal and distal cells.

Most studies in the past decades have focused on a limited number of communication molecules in a given anatomical site or physiological process. The availability of large-scale transcriptomic datasets from several cell types, tissue locations, and cell activation states, opened the possibility of reconstructing cell-to-cell interactions based on the expression of specific ligand-receptor pairs on sender and target cells, respectively. Many of them exploit single cell RNAseq datasets to infer communication between groups of cells within the same dataset^4–7^. Despite leading to interesting and often innovative hypotheses^4,6,8^, these methods do not integrate putative signals that may come from more distant cells. Also, they cannot be applied to bulk transcriptomic data derived from a given cell population. Such datasets are numerous in public databases, and can be a source of novel insights into how each cell type may send or receive communication signals.

Another important aspect when inferring cell-to-cell communication is the use of databases of ligand-receptor interactions. Some are very broad with over 2000 ligand-receptor pairs^9^, but lack systematic manual or expert curation, which may impact the quality and biological relevance of the annotation. Others include lower numbers of ligand-receptor pairs and provide manually curated information from the literature^4,10^, without necessarily providing systematic combinatorial rules for the association of protein subunits into multimeric ligands or receptors.

The last point relates to the granularity that is structuring the biological information into families and subfamilies of functionally and structurally related molecules. We only found one tool that provides a classification into four families of communication molecules^5^, while suffering from other limitations in particular the lack of manual curation.

In this study we developed ICELLNET, a novel and versatile computational framework to infer cell-to-cell communication from a wide range of bulk and single cell transcriptomic datasets. Each family of communication molecules was expert curated and organized into biologically relevant sub-families. ICELLNET offers an array of visualization tools in order to facilitate biological interpretation and discoveries. We provide applications to public datasets, and our own original transcriptomic datasets in non-immune (tumor fibroblasts) and immune cell types. Experimental validation of ICELLNET-derived predictions demonstrated IL-10 control of human dendritic cell communication.

## Results

### Expert-curated database of ligand-receptor interactions

In order to globally reconstruct cell communication networks, we curated a comprehensive database of ligand-receptor interactions from the literature^3,11,12^ and public databases^10,13^. Rather than focusing on the breadth of the resource, we used a strategy that prioritizes expert manual curation and biological insight based on precise biochemical and functional classifications. This led to the integration of 380 ligand-receptor interactions into the ICELLNET database (**Suppl. Table S1**). Whenever relevant, we took into account the multiple subunits of the ligands and the receptors (**Figure 1A**). Interactions were classified into 6 major families of communication molecules, with a strong emphasis on inflammatory and immune processes: Growth factors, Cytokines, Chemokines, Immune Checkpoints, Notch signaling and Antigen binding (**Figure 1B and Suppl. Table S1**). Other families such as hormones or adhesion molecules were more scarcely represented. In order to simplify the subsequent graphical visualization, these were grouped as “other” in our current classification (**Figure 1B and Suppl. Table S1)**.

**Fig. 1:**
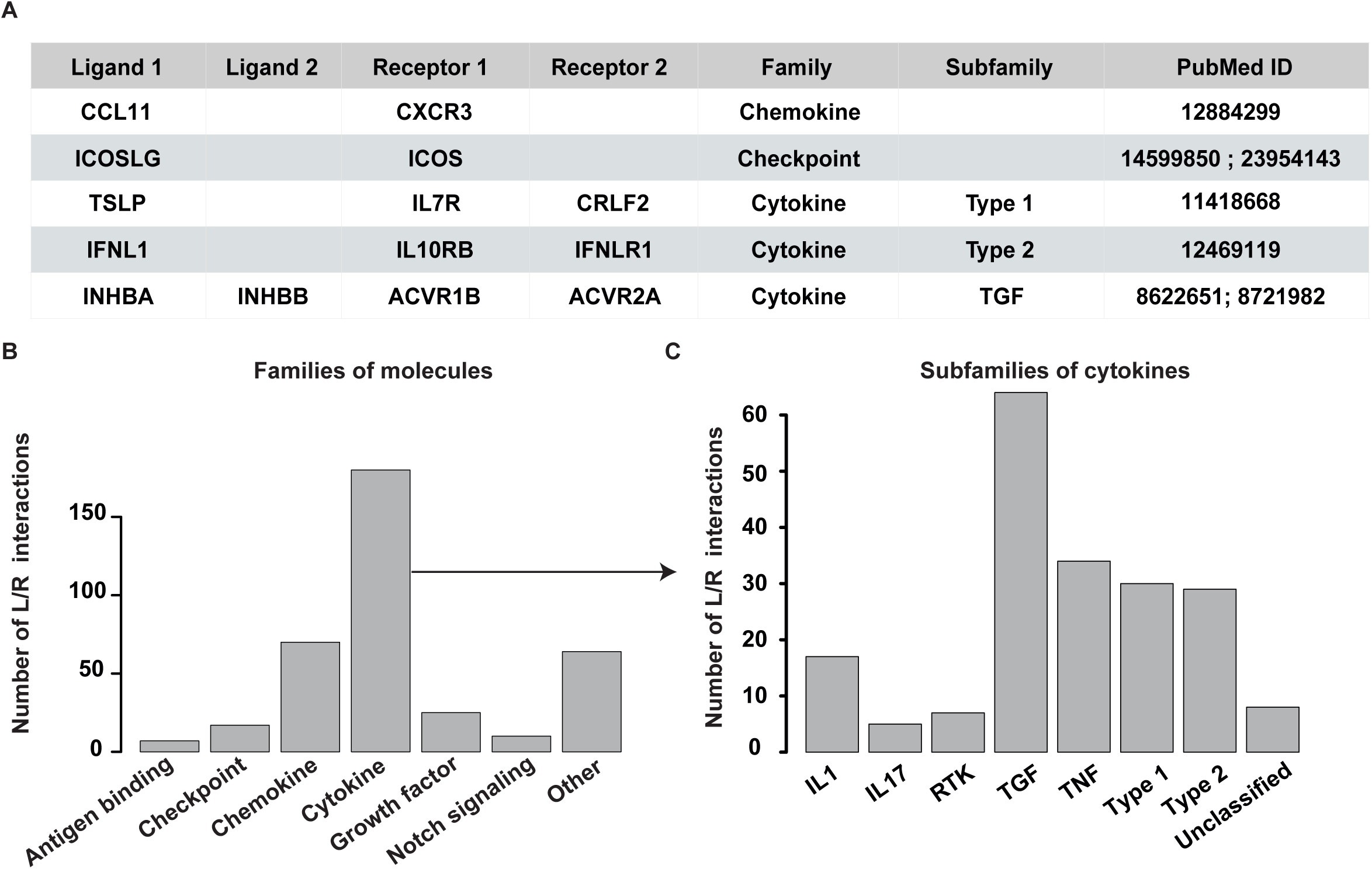
Structure of the ligand-receptor database. **(A)** Extract of the ligand-receptor database **(B)** Histogram of the number of communication interaction by families of molecules, **(C)** Histogram of the number of communication interaction by subfamilies of cytokines.

Cytokine-receptor pairs were mapped in an exhaustive manner, by exploiting a series of reference articles and consensus classifications. They represent 50% of the total interactions included in the database (194 interactions), and were further classified into 7 sub-families according to structural protein motifs: type 1 cytokines, type 2 cytokines, IL-1 family, IL-17 family, TNF family, TGF-ß family and RTK cytokines ^3,14–16^(**Figure 1C**).

### Development of a computational pipeline to dissect intercellular communication

In the ICELLNET framework, we developed an automatized tool in R script to infer communication between multiple cell types by integrating; 1) prior knowledge on ligand-receptor interactions **(Figure 1)**; 2) computation of a communication score between pairs of cells based on their transcriptomic profiles, and; 3) several visualization modes to guide results interpretation. Quantification of intercellular communication was achieved by scoring the intensity of each ligand-receptor interaction between two cell types from their expression profiles **(Figure 2)**.

**Fig. 2:**
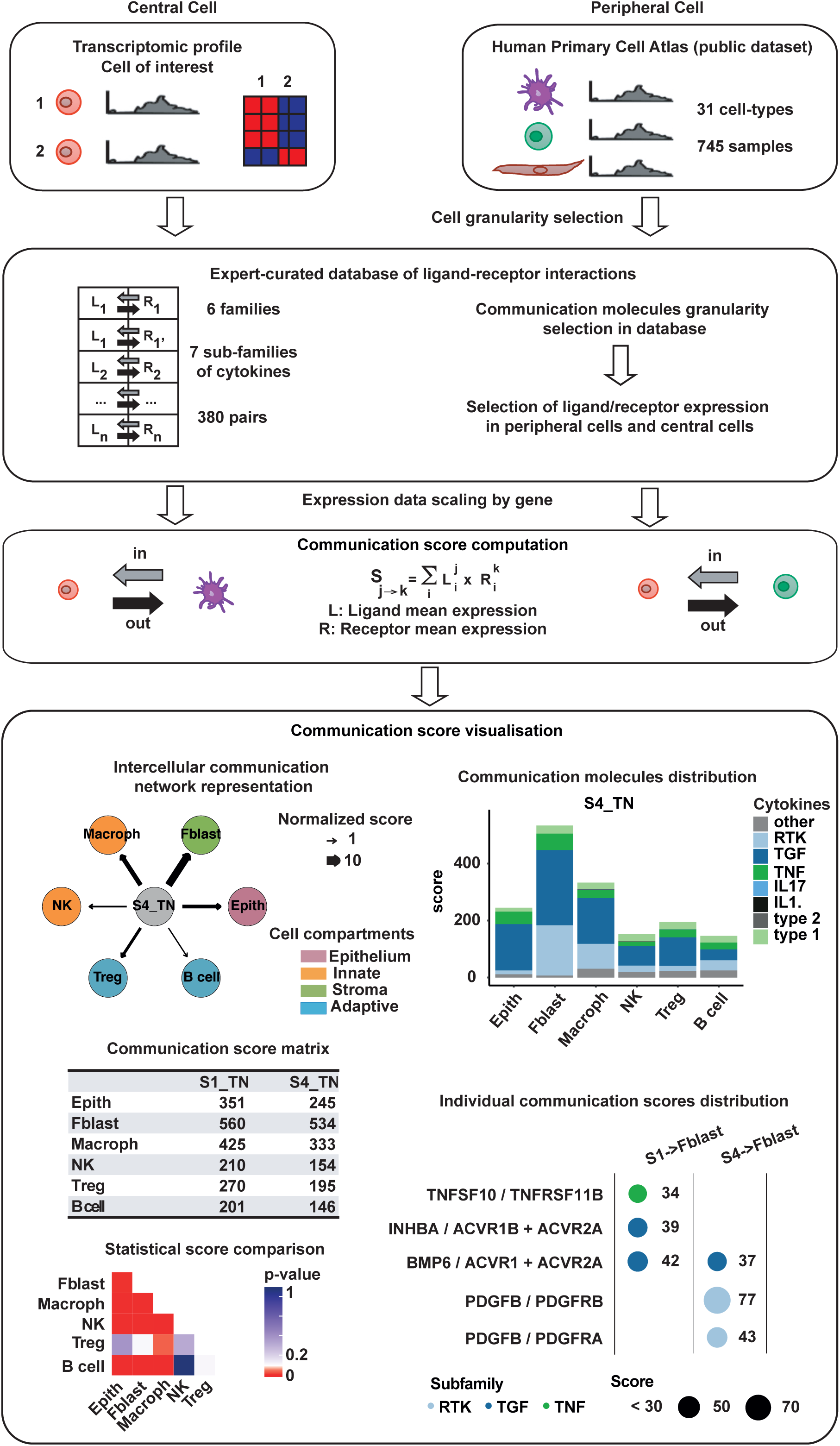
ICELLNET pipeline to study intercellular communication from cell transcriptional profiles. Pipeline used to create the intercellular communication score and network reconstruction.

From each transcriptomic profile, all genes or only differentially expressed genes could be used, and no filtering threshold was applied to gene expression. Taking advantage of the ICELLNET database, the genes coding for ligands/receptors were selected from all 380 interactions to compute the score, but it is also possible to restrict the database to specific families of molecules, depending on the biological question.

A unique feature and strength of ICELLNET is its ability to infer cell-to-cell communication even from an individual cell population-based transcriptome of interest (the “central cell”). ICELLNET first considers the transcriptomic profile of each “central cell”, which may correspond to different cell subsets or the same cell type cultured in different biological conditions (**Fig 2 top-left)**. ICELLNET separately considers other cell types with known transcriptomic profiles that can connect to the central cell, called « peripheral cells ». These can be cell types coming from the same dataset as the central cell, or from any other transcriptomic dataset. The ICELLNET pipeline has integrated reference transcriptomic profiles by using the Human Primary Cell Atlas^17,18^. This public dataset includes transcriptomic profiles of 31 cell types including immune cells, stromal cells, neural cells, and tissue specific cell types, all generated with the same Affymetrix technology **(Fig 2 top-right)**. Human Primary Cell Atlas has been downloaded and added to ICELLNET framework, in order to be used as reference transcriptomic profiles of peripheral cell types **(Suppl Table S2)**.

### Establishment of a score to assess the communication between cells

From the transcriptomic profiles, we selected the genes coding for the ligands and the receptors in our database. We designed the tool to enable the user to focus either on the ligands and/or receptors that are differentially expressed between conditions of study, or to use the entire ligand-receptor database to compute the communication score.

Since cell-to-cell communication is directional, we considered ligand expression from the central cell, and receptor expression from the peripheral cells in order to assess outward communication. Conversely, we then selected receptor expression from the central cell, and ligand expression from peripheral cells in order to assess inward communication (**Figure 2 middle**). Gene expression levels were scaled to avoid a communication score driven predominantly by highly expressed genes. In the ICELLNET framework, quantification of intercellular communication consists of scoring the intensity of each ligand-receptor interaction between two cell types with known expression profiles. Whenever relevant, we took into account multiple ligand subunits, or receptor chains, using logical rules to impose their co-expression in order to consider functionality. The score of an individual ligand-receptor interaction was computed as the product of their expression levels by the respective source (central) and target (peripheral) cell. When a communication molecule (ligand or receptor or both) was not expressed by a cell, the score of this particular interaction was set to zero. Individual scores were then combined into a global metric assessing the overall exchange of information between the cell types of interest (**Figure 2 middle**), defining a global communication score. ICELLNET provides a matrix summarizing all global communication scores as an output of these analytical steps.

### ICELLNET offers different graphical representations allowing multiple layers of interpretation

ICELLNET generates a large quantity of data and scores, which are complex to interpret and analyse. In order to facilitate hypothesis generation, three graphical representations were generated to help visualise and interpret the results (**Figure 2 bottom**). The first representation allows the visualization of intercellular communication networks in directed connectivity maps. In these graphs, nodes represent cell types, the width of the edges connecting two cell types is proportional to their global communication score and the arrows indicate the direction of communication. The second visualization mode breaks down the global scores into the contribution of specific molecular families through a barplot representation. This allows the identification of patterns of co-expressed molecules from the same family, potentially contributing to coordinated biological functions. We implemented statistical analyses of the scores (see Methods) to evaluate the robustness of the differences between scores. The resulting p-values can be visualized as an additional heatmap. The third representation displays the highest contributing ligand-receptor pairs to the communication score within a given channel in a balloon plot. This enables the identification of specific interactions that may drive the global intercellular communication. Thus, the ICELLNET framework is a powerful tool to assess intercellular communication with different visualisation modes that can be helpful to dissect underlying mechanisms and extend biological knowledge and understanding.

### Application of ICELLNET to study human breast cancer-associated fibroblasts

Cancer-associated fibroblasts (CAFs) are stromal cells localized in the tumor microenvironment that are known to enhance tumor phenotypes, notably cancer cell proliferation, and inflammation. Recently, four subsets of CAFs have been identified and characterized in the context of previously untreated Luminal and Triple Negative Breast Cancer (TNBC)^19^. CAF-S1 and CAF-S4 specifically accumulated in TNBC and CAF-S1 was associated with an immunosuppressive microenvironment. This study raised important questions about the regulatory mechanisms involved, in particular the role of cell-to-cell communication. Using the available transcriptional profiles of CAF-S1 and CAF-S4 in TNBC (**Figure 3A**), we applied the ICELLNET pipeline to reconstruct the intercellular communication network with 14 other cell types potentially localized in the tumor microenvironment (TME) (**Figure 3B and Suppl. Table S3A-B**). The peripheral cells were selected from Human Primary Cell Atlas and included innate immune cells (monocytes, macrophages, pDC, DC1, DC2, NK cells, neutrophils), adaptive immune cells (CD4+ T cells, CD8+ T cells, Tregs, B cells), epithelial and stomal cells (fibroblasts and endothelial cells). In order to assess the global intercellular communication, we first used the network graphical vizualisation. This strongly suggested that CAF-S1 has a greater communication potential than CAF-S4 (**Figure 3B**). The rescaled communication scores were higher for CAF-S1 compared to CAF-S4, and the differences were statistically significant for epithelial cells (score CAF S1 > Epith = 6, score CAF S4 > Epith = 4, pvalue< 0.1), endothelial cells (score CAF S1 > Endoth = 6, score CAF S4 > Endoth = 4, pvalue < 0.1), plasmocytoid dendritic cells (score CAF S1 > pDC = 6, score CAF S4 > pDC = 4, pvalue < 0.1) and B cells (score CAF S1 > B cells = 3, score CAF S4 > B cells= 1, pvalue < 0.1) (**Figure 3B, 3C and Suppl. Table S3A-B**).

**Fig. 3:**
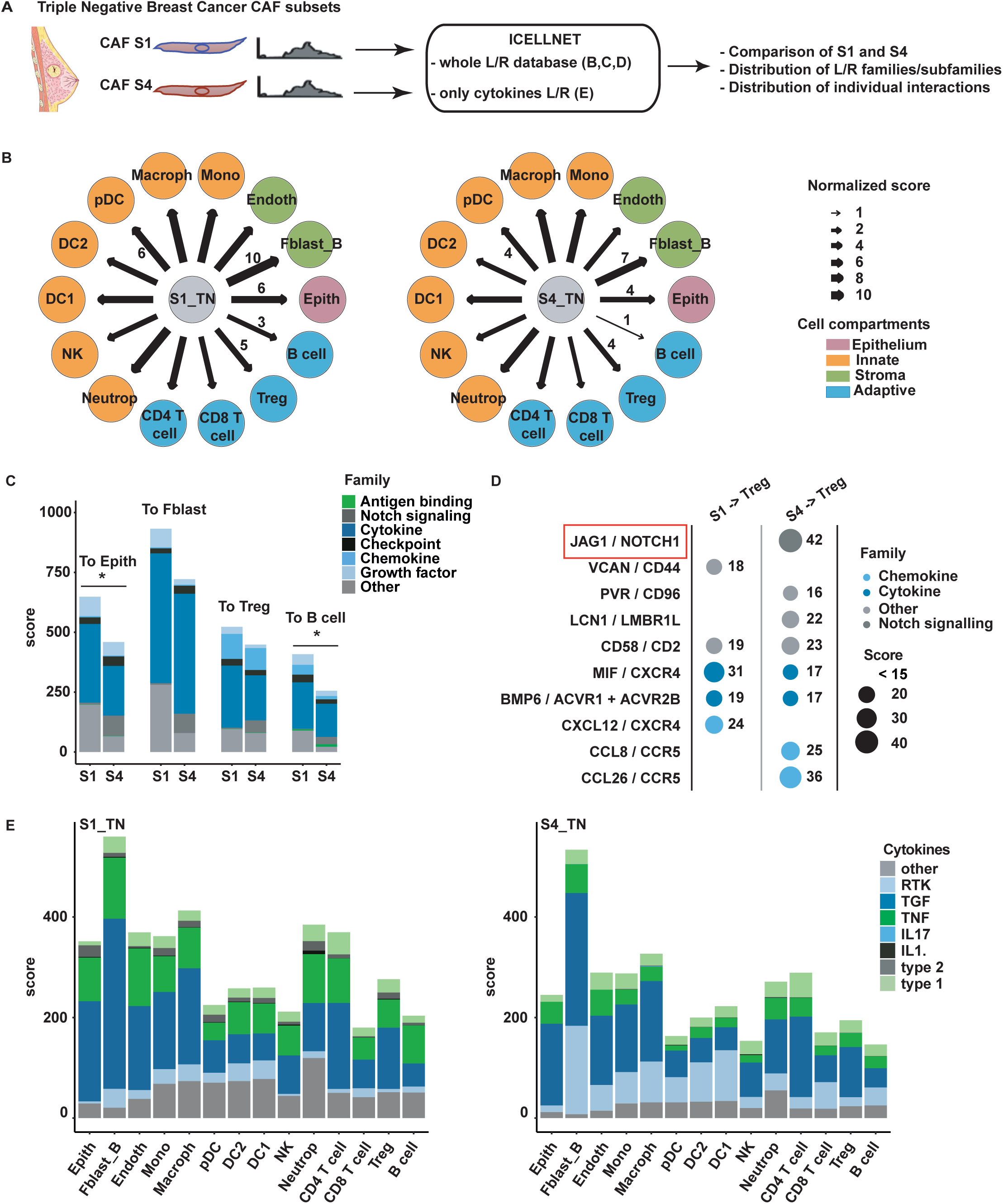
Dissecting intercellular communication between Triple-Negative breast cancer infiltrating CAF subsets. **(A)** Workflow of the analysis. **(B)** Connectivity maps describing outward communication from CAF-S1 (n=6) and CAF-S4 (n=3) subsets to primary cells. The CAF subsets are considered as central cells and colored in grey. Primary cells are considered as peripheral cells and are colored depending on the cell compartment (green: stroma, orange: innate, blue: adaptive, pink: epithelium). The width of the edges corresponds to a global score combining the intensity of all the individual ligand/receptor interactions. A scale ranging from 1 to 10, corresponding to minimum and maximum communication scores, is shown in the legend. A selection of normalized scores is written directly on the network. **(C)** Barplot of communication score with contribution by families of communication molecules between CAF subsets and a selection of peripheral cells. Significant differences are shown on the graph (*: p-value ≤ 0.1). **(D)** Balloon plot of individual interaction scores between CAF subsets and Tregs. Two biologically-interesting communication channels were highlighted by red boxes. **(E)** Barplot of communication score with contribution restricted to cytokines subfamilies between CAF subsets and a selection of peripheral cells.

### CAF-S4 uses specific communication channels to interact with the TME components

We focused on the biological composition of the score, to identify families of molecules highly involved in CAFs communication with the selected cells. We selected 4 peripheral cell types: epithelial cells, fibroblasts, Tregs and B cells. Using the barplot representation, we looked for differences between CAF-S1 and CAF-S4 in terms of genes coding for families of communication molecules. We found that genes coding for communication molecules inducing Notch signaling were specifically expressed by CAF-S4 to communicate with other cells (**Figure 3C**). Looking at individual communication interactions between CAF subsets and Tregs demonstrated that gene coding for JAG1 protein was only expressed by CAF-S4 to interact with NOTCH receptors (NOTCH1 and NOTCH2 genes expressed), and thus potentially having a role in activating the Notch signaling pathway (**Figure 3D and Suppl. Table S3A-B**). For both CAF subsets, the barplot representation indicated that cytokines-receptors interactions were highly contributing to the global communication scores compared to other families of molecules (**Figure 3C**). This observation led us to focus on cytokine-mediated communication using the ICELLNET pipeline (**Figure 3E**). By considering only cytokine-receptor interactions, the CAFs appear to communicate more with other fibroblasts compared to other cell types with a significant p-value (**Figure 3E, Suppl. Figure S1A**). Also, this approach highlighted that RTK cytokines, and notably PDGFB coding for PDGF, were preferentially expressed by CAF-S4 compared to CAF-S1 (**Figure 3E, Suppl. Figure S1B and Suppl. Table S3C**). We also applied ICELLNET pipeline to study inward communication between the peripheral cells and the CAF subsets, which revealed no difference between CAF-S1 and CAF-S4 in term of communication score intensities but also in terms of the families of molecules involved in communication (**Suppl. Figure S2**). Thus, the ICELLNET framework allowed us to identify specific communication channels revealing potential interactions between CAF-S4 and TME components.

### Application of ICELLNET to study communication between specific immune cells

After using TNBC-infiltrating CAFs dataset to test the connectivity map reconstruction, we wanted to test the hypothesis that the ICELLNET tool would allow us to characterize cellular communication using the immune system as a model. Particularly, we were interested in studying communication of resting and perturbed immune cells. To explore the role of autocrine loops, we cultured LPS-activated human monocyte-derived dendritic cells (DCs) in the presence or absence of blocking antibodies (Abs) to the TNF and IL-10 receptors (aTNFR and aIL-10R). No effect on cell viability was observed (**Suppl. Figure S3A**). The most prominent effect of LPS on DC hallmark maturation markers was observed at the mRNA level in the time frame of 4 to 8 hours following activation^20^. We performed large-scale microarray analysis after 4 and 8 hours of culture of DC with LPS, with and without blocking Abs to TNF and IL-10 receptors (**Figure 4A**).

**Fig. 4:**
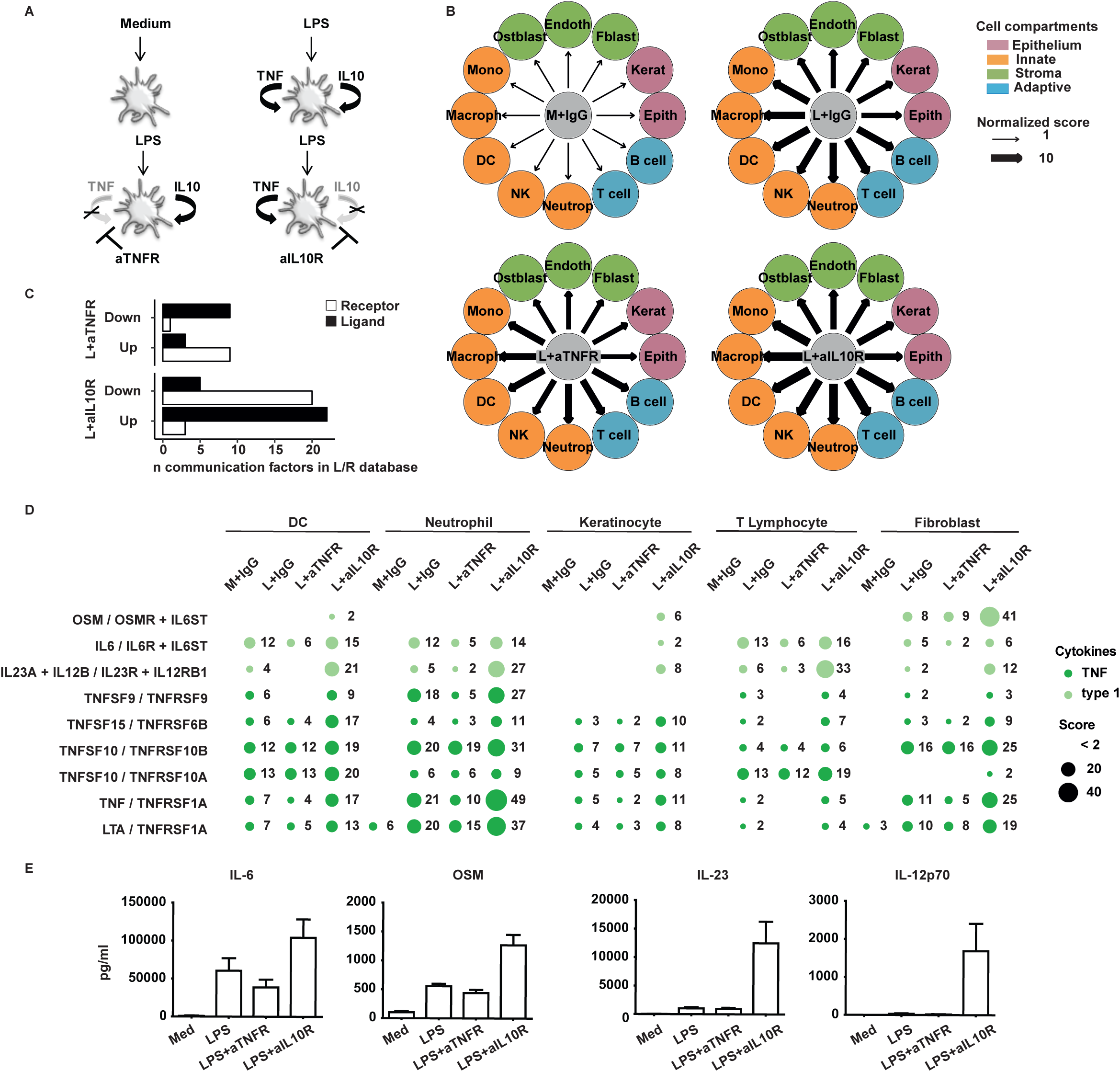
IL-10R blocking activates a cell-to-cell communication module in LPS-stimulated DCs. **(A)** Depicted are the 4 experimental conditions for which transcriptomics was generated (n = 6). **(B)** Connectivity maps describing outward communication from DCs to putative target cells in the conditions: Med, LPS, LPS+αTNFR and LPS+αIL-10R. Twelve primary cell types are considered as peripheral cells and are colored depending on the cell compartment (green: stroma, orange: innate, blue: adaptive, pink: epithelium). The width of the edges corresponds to a global score combining the intensity of all the individual ligand/receptor interactions, normalized to the medium condition. A scale ranging from 1 to 10, corresponding to minimum and maximum communication scores, is shown in the legend. **(C)** Gene corresponding to ligands (black) and receptors (white) counted in each loop signature and plotted according to regulation directionality: upregulated (Up) or downregulated (Down). Genes with separability score ≥ 4 were included in each condition’s signature. (**D)** Protein levels of IL-6, OSM, IL-23 and IL-12p70 (means ± SEM), demonstrating increased secretion in LPS+αIL-10R DC supernatant.

Despite extensive studies of both TNF and IL-10 in the context of innate immunity, their different contribution to DC intercellular communication could not be predicted *a priori* at this systems level. We applied ICELLNET to reconstruct the intercellular networks between DCs and the putative target cells. The network representation demonstrated an increase of the global communication score in all 12 channels, when comparing 8-hours LPS-activated DC to resting (medium) DC (**Figure 4B)**. Importantly, these maps revealed that blocking the IL-10 loop determined the largest amplification of DC communication with all 12 cellular targets, while the blocking of TNF loop in LPS-activated DCs had a negligible effect on the global communication score (**Figure 4B, Suppl. Figure S3B and Suppl. Table 4A-E)**

### IL-10 controls an intercellular communication module in LPS-activated dendritic cells

We compared the transcriptomic profiles of each condition (aTNFR and aIL-10R) to the LPS-alone condition to extract the differentially expressed genes (DEG) (**Suppl. Table 4F**). We then screened the IL-10 and TNF DEG to identify ligands and receptors included in the database. We were able to extract 27 ligands and 23 receptors which were differentially regulated from the aIL-10R condition, while there were only 12 ligands and 10 receptors differentially regulated from the aTNFR condition (**Figure 4C**).

ICELLNET barplots suggested that cytokines were driving the increase in the communication score when blocking IL-10R. We looked at the subfamilies of cytokines to precisely identify the key communication channels (**Suppl. Figure S3B**). Type 1 and TNF subfamilies were increased in aIL-10R condition compared to others. This was confirmed by individual channel communication scores (**Figure 4D**). To confirm the hypothesis that IL-10 controls cytokine-mediated DC communication, we selected four important immunoregulatory molecules from the IL-6- and IL-12-families, and further validated expression at the protein level in 24 hours culture DC supernatants using cytometric bead array (CBA) and ELISA (**Figure 4E**).

### Experimental validation of multiple IL-10-dependent communication channels

To assess communication efficiency, i.e. how increased connectivity translates into functional changes in target cells, we turned to experimental validation of predicted communication channels using immunological assays adapted to the output response of each cell type. Due to its physiopathological relevance, we first investigated the DC-T cell axis through co-culture experiments of T cells with DCs treated by LPS with or without TNFR and IL-10R blocking antibodies (**Figure 5**). We found that naive CD4+ T cells, when co-cultured with LPS-DC in the absence of the IL-10 loop, globally increased and shifted their pattern of cytokine secretion, compared to LPS-DCs, while blocking the TNF loop had almost no effect (**Figure 5A**). Similar results were obtained with memory T cells (**Figure 5B**).

**Fig. 5:**
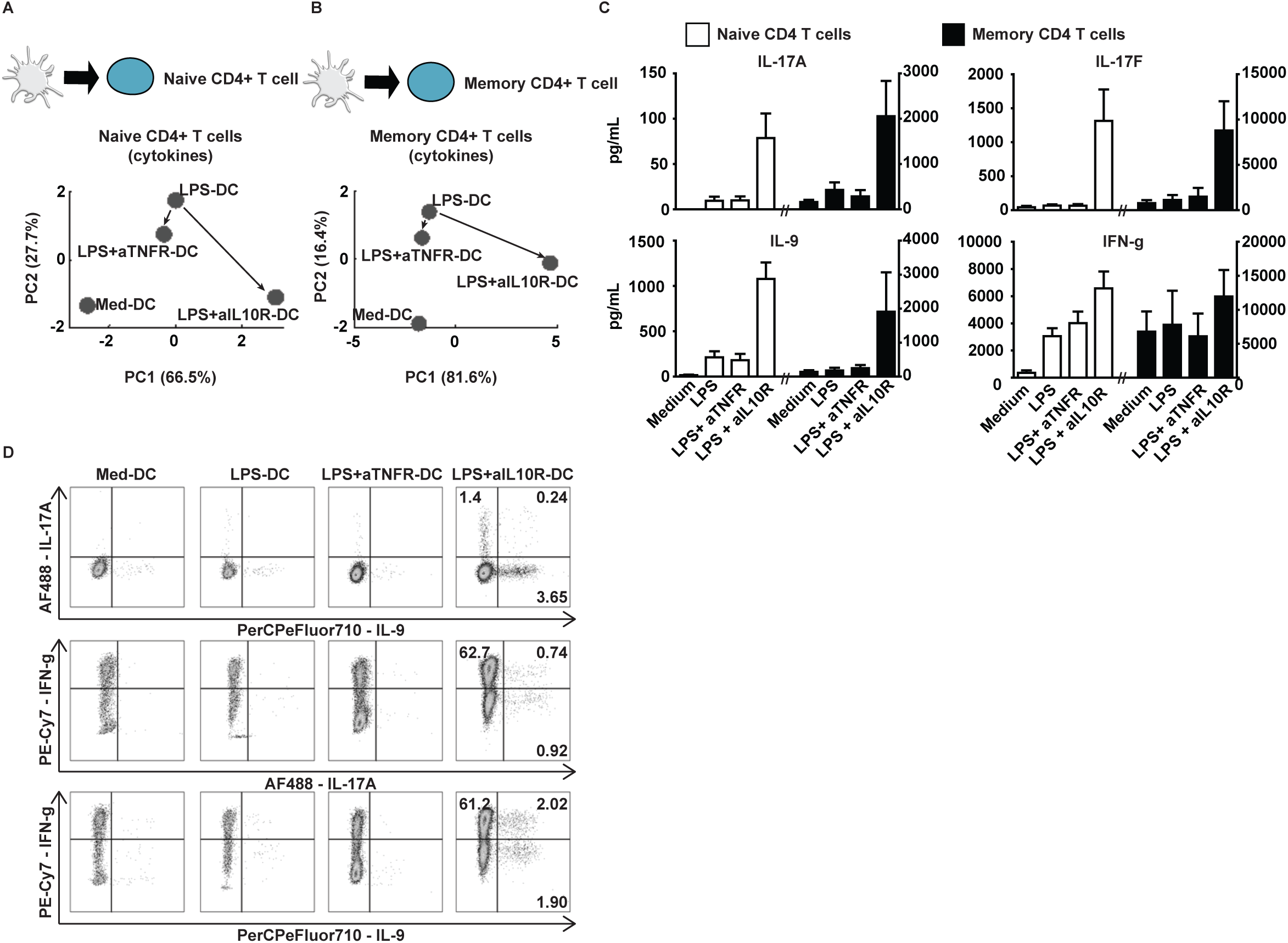
IL-10 but not TNF loop dictates T helper polarization by LPS-DC. **(A-B)** Supernatants of CD4+ naive (A) and memory (B) T cells, co-cultured with the indicated DCs, were analyzed for the presence of T helper cytokines by CBA: IL-2, IL-3, IL-4, IL-9, IL-10, IL-17A, IL-17F and IFN-γ (A) and all the above in addition to IL-5, IL-13 TNF and GM-CSF (B). Results are shown in a two-dimensional principal component analysis (PCA). Dots represent mean of 9 (A) or 6 (B) independent co-culture experiments. **(C)** Histogram representation (means ± SEM, n = 16) of 4 cytokines present in the supernatant of naive (white bars, left axis) or memory (black bars, right axis) supernatant. **(D)** CD4+ naive T cells were analyzed for IL-17A, IL-9 and IFNg production using intracellular staining FACS. Percentage of positive producers is given. Shown is one representative out of 3 independent experiments.

Since the IL-10/IL-10R pathway may have a direct effect on T helper cells during the differentiation process, we verified that the observed T helper polarization was indeed due to the IL-10 loop blockade in the DCs, and not due to a direct effect on T cells (**Suppl. Figure S4A**). It is possible that residual IL-10R blocking antibodies could have acted directly on T cells during DC-T co-culture. By adding IL-10R antibodies during DC-T co-culture (not only during DC activation) we demonstrated that any IL-10R antibodies in this setting would not have any direct effect on T cell polarization.

Among the factors explaining the secretion profile of T cells determined by LPS+aIL-10R-DCs, we observed a remarkable emergence of Th17 cytokines (**Figure 5C**), in line with recent murine studies^21,22^. Strikingly, IL-9 secretion was also increased (**Figure 5C**), and produced by a T cell population distinct from the Th17 cells producing IL-17A alone or co-expressed with IL-9 and IFN-g (**Figure 5D**). This provides the first demonstration that LPS-activated DCs, in the absence of an IL-10 loop, determine Th17 and Th9 polarization in humans, both of which participate in host defense and autoimmunity^23,24^.

In order to validate the model-based hypothesis that there is increased communication between DC and multiple cell types, we considered three additional types of target cells: keratinocytes, plasmacytoid DCs (pDC) and neutrophils. Similar to T cells, these cell types play key roles in the inflammatory microenvironment and had an increased global communication score. Target cells were cultured with DC-derived supernatants, and their activation assessed by qRT-PCR or FACS. LPS-DC supernatant induced marginal keratinocyte activation, as assessed by the expression of TNF, IL-1β and this was not affected by aTNFR (**Figure 6A**). However, blocking the IL-10 loop dramatically increased both factors (**Figure 6A**), validating a potent DC to keratinocyte communication controlled by IL-10. This extends DC-induced keratinocyte activation^25,26^ to the context of bacterial infection.

**Fig. 6:**
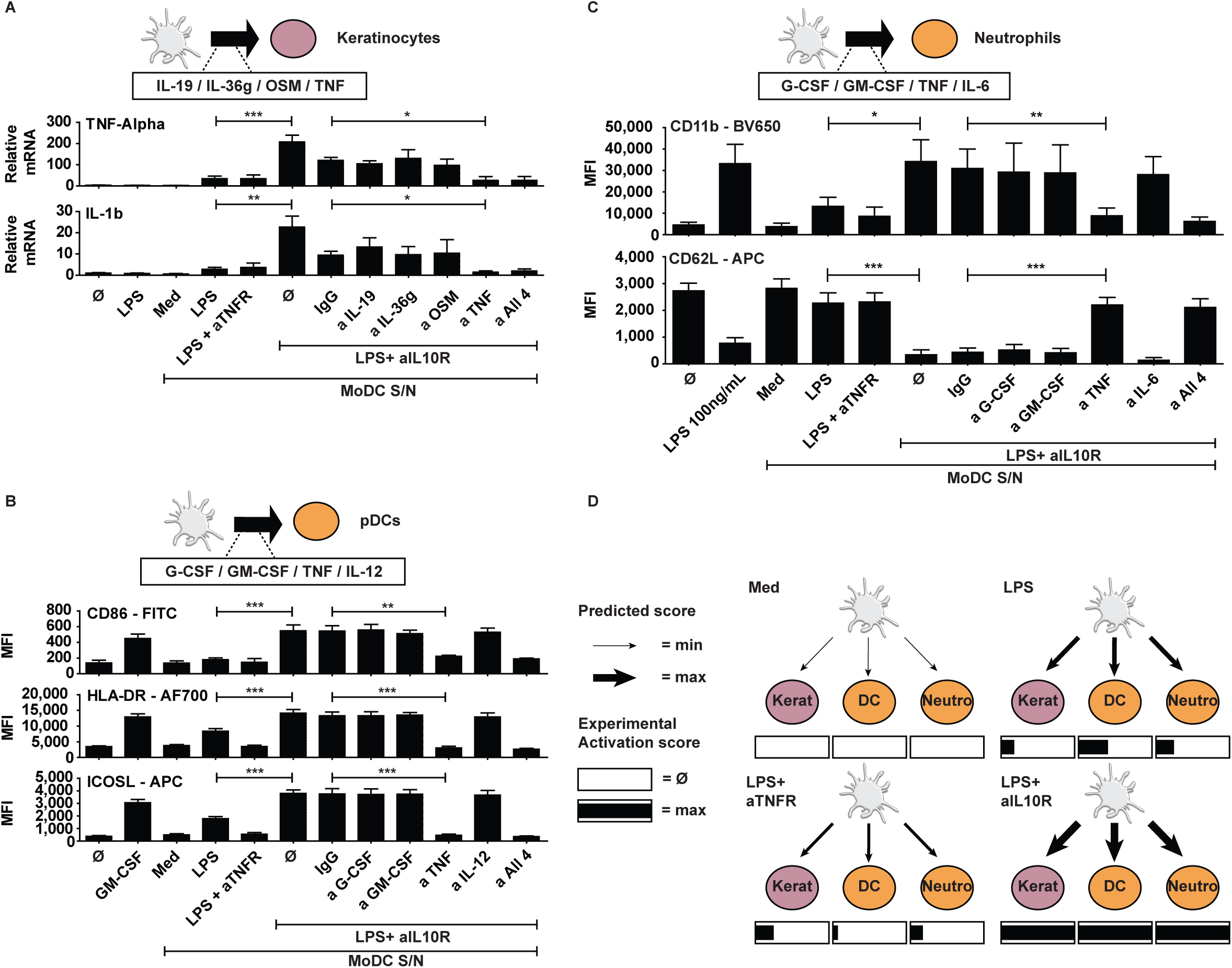
IL-10 loop controls DC communication with keratinocytes, neutrophils and pDCs. **(A)** RT-PCR analysis of the expression of TNF and IL-1b mRNA in HaCat cells incubated with medium, LPS or with supernatant (diluted 1:10) of the indicated DCs for 4 hours. Blocking antibodies for the cytokines IL-19, IL-36g, OSM and TNF were added to LPS+αIL-10R-DC supernatant for 1 hour incubation before culturing with HaCat cells. Data represent mean ± SEM, n=4, * *p*<0.05. **(B-C)** Expression of maturation markers CD86, HLA-DR and ICOSL (B) or DC11b and CD62L (C) analyzed by flow cytometry with surface staining on pDCs (n=18) cultured with supernatant (diluted 1:10) of the indicated DC for 24 hours (b) and neutrophils (n=9) cultured with supernatant (diluted 1:100) of the indicated DC for 1h. Blocking antibodies for the cytokines GCSF, GM-CSF, TNF and IL-12 (for pDC) or IL-6 (neutrophils) were added to LPS+αIL-10R-DC supernatant for 1 hour incubation before culture. Each biological replicate comprised independent DC donor paired to independent pDCs / neutrophils donor. Data represent mean ± SEM, * *p*<0.05; ** *p*<0.01; *** p<0.001 (paired *t*-test). **(D)** For each target cell, we reduced the different activation markers to a single parameter normalized between 0 (Ø) and 1 (max) in the rectangles. The value 0 corresponds to the activation level induced by supernatants from untreated DC, while 1 corresponds to the maximum activation level from all the observed conditions. These experimentally validated activation scores were in qualitative agreement with the model predictive intensity scores of communication between DC and the target cells, represented by the width of the edges.

The DC-pDC communication channel was also controlled by IL-10, since LPS+aIL-10R-DC supernatants activated pDCs (as assessed by CD86, HLA-DR, and ICOSL surface expression), in comparison to LPS-DCs (**Figure 6B**). DC-induced activation of pDC and keratinocytes was not due to the presence of residual aIL-10R (**Suppl. Figure S4B and C**). DC-pDC crosstalk was suggested to be important in antiviral^27^, antibacterial^28^, and antitumor^29^ immune responses. Through our systems approach, we have shown that IL-10 controls DC-pDC connectivity.

Neutrophils contribute to DC migration to infection sites and to their subsequent activation^30,31^. Reciprocally, it was proposed that DCs can promote neutrophil survival^32^. LPS-DC supernatant induced only a mild activation of neutrophils (as evaluated by rapid upregulation of CD11b with concomitant downregulation of CD62L), while LPS+aIL-10R-DC supernatants led to a strong activation of neutrophils (**Figure 6C**), establishing an IL-10 loop control of DC-neutrophils communication.

For all the above-mentioned communication channels, we aimed at getting further mechanistic insight. First, we performed control experiments using exogenous LPS that formally excluded any direct effect of LPS at the concentrations found in the DC supernatants (**Suppl. Figure S4D**). We then considered ligand-receptor interactions showing high intensity, and thus more likely to mediate cellular crosstalk as observed with the LPS+αIL-10R-DC supernatants (**Suppl. Table S5**). We blocked, in each DC communication channel, 4 of the ligands known as potential activators of the target cell type: GCSF, GM-CSF, IL-6 and TNF for neutrophils, IL-19, IL-36 gamma, OSM and TNF for keratinocytes, and G-CSF, GM-CSF, TNF and IL-12 for pDCs. Importantly, blocking TNF alone in the LPS+αIL-10R-DC supernatant was sufficient to inhibit keratinocyte, pDC and neutrophil activation (**Figure 6A-C**). By comparing the predicted communication intensities with a global score describing the activation level of keratinocytes, pDC and neutrophils, we observed a qualitative agreement (**Figure 6D**), demonstrating increased communication efficiency. In all cases, the target cells were most activated in LPS+aIL10R condition.

## Discussion

The majority of studies which aim to reconstruct intercellular communication from transcriptomic datasets integrate prior knowledge in the form of a ligand-receptor interaction database. This provides a straightforward manner to infer communication when a match is identified between a ligand and a cognate receptor for two respective cell types. The largest of such databases^9^ integrated over 2500 ligand-receptor pairs through literature mining and computational analysis, and has been exploited in multiple computational tools for predicting cell-to-cell communication^5,8,33,34^. However, this approach lacks experimental validation of predicted ligand-receptor interactions, and it does not take into account the different subunits of ligands or receptors. With ICELLNET, we have developed a fully manually curated database, combining biological relevance, ease of use, and experimental validation. Except for one study^5^, ICELLNET is the only database offering a classification of predicted interactions into biological families. Similar to CellPhoneDB^10^, ICELLNET takes into account the multiple subunits of ligands and receptors, by introducing logical rules for co-expression of protein subunits. A systematic comparison of cytokine interactions revealed 14 interactions included in ICELLNET but not in CellPhoneDB, such as MIF/CXCR2 and MIF/CXCR4^35^. Although ICELLNET includes a relatively small number of interactions compared to other existing databases, it is very specific and exhaustive for cytokine interactions, and will in time be extended to all chemokine and checkpoint interactions, thus providing a unique resource to study intercellular communication within the immune system.

To make ICELLNET a valuable resource, we have established a strategy to keep the database updated and integrate missing knowledge. A significant number of interactions have been established in the past 20 years, but there are still receptors without known ligands, such as TNFRSF21 (DR6), RELT, TROY and NGFR from TNF receptor family^36^, and ligands without known receptors such as IL17D^16^. New receptors for existing ligand-receptor pairs can also be uncovered using this approach. For example, even though it was already known that IL34 and M-CSF could separately activate M-CSFR^37^ it was then described that IL34/M-CSF heterodimer was also capable of activating M-CSFR^38^. We will apply a PubMed alert strategy to cover all new interactions that could be described on these pre-identified ligand and/or receptor partners.

Existing tools infer communication between cells from scRNAseq datasets^4,5,7^. We have designed ICELLNET as a versatile tool, which can be applied to bulk cell profiles (Affymetrix or RNAseq) widely available in public databases, but also to fully documented scRNAseq datasets to infer communication between clusters or groups of cells. This can be easily adapted to other types of data such as flow cytometry data. By using the Human Primary Cell Atlas as a reference for transcriptomic profiles^17,18^, ICELLNET allows us to integrate cell communication partners (sender or receiver) not included in a given original experimental dataset. We identified the Human Primary Cell Atlas as a particularly suitable resource, as it integrates transcriptional profiles of over thirty human primary cell types generated with the same Affymetrix platform^18^. While previous applications of this atlas enabled the identification of specific tissue-related genes^39,40^, we developed an original use for this resource to simulate cell cross-talks in diverse microenvironments. In addition, ICELLNET can accommodate other original RNAseq datasets of cell populations^41,42^ as reference profiles to infer intercellular communication. Hence, ICELLNET is an extremely flexible tool, which can be easily adapted depending on the biological question, by offering the possibility to select communication molecules families and cell types of interest.

A key aim in studying cell-cell communication is to represent cellular interactions in a clear and biologically relevant manner. Visualization is important to understand the different levels of interactions, at the cellular and molecular levels. Most of the available tools use two main graphical representations; heatmaps and circos plots. These complex plots represent all possible interactions at once and can be difficult to read and interpret. ICELLNET offers four original visualization modes with different properties to represent cell-to-cell communication from a global view of specific interactions. These different representations simplify interpretation of the results, help users to elaborate hypotheses and allow in-depth analysis of cell-to-cell interactions.

The cytokine family of communication molecules plays a key role in homeostatic processes, such as cell development and differentiation, tissue homeostasis, and inflammation^3,14,43^. In the past 20 years, a large number of new cytokines have been identified, cloned, and studied to elucidate their biological function. This has significantly enriched the classification of cytokines into structural families matching evolution and functional processes^3,44^. ICELLNET is now providing an exhaustive and expert curated resource of all known cytokines and their receptor interactions, according to reference knowledge. This opens possibilities for researchers to decipher complex cytokine-mediated communication, and the implication of specific cytokines in disease.

Fibroblasts are important structural stromal cells at steady state and inflammation. Yet, how they communicate with neighboring cells is not well described. Applying ICELLNET to breast cancer fibroblasts’ bulk cell transcripts revealed potentially novel interactions between CAF subsets and tumor microenvironment components. The CXCL12/CXCR4 interaction that we found within CAF-S1-to-Tregs (Fig 3D) was also described in other studies^19,45^, and contributes to the immunosuppressive phenotype displayed by CAFS1. ICELLNET also highlighted interactions specific to CAF-S4 subset such as JAG1 with Notch receptors (NOTCH1, NOTCH2), and expression of PDGF proteins interacting with their cognate receptor. These proteins have never been associated specifically to CAF-S4 subset at the transcriptomic level and warrant further experimental validation studies.

Other studies have shown that IL-10 regulates DC-derived inflammatory cytokines and chemokines, in particular IL-12^46^, and that IL-10 secreted by LPS-activated DCs controls a communication channel in an autocrine manner^47^. Through our systems approach, we could demonstrate that endogenous DC-derived IL-10 governs the global connectivity of DCs with multiple cell types. This original *in vitro* dataset also provides an experimental validation of our intercellular communication hypotheses, making ICELLNET the first experimentally validated tool to assess intercellular communication.

Thus, ICELLNET is an adaptable tool that allows us to gain insight into communication channels between cells from one bulk transcriptomic profile of a cell population. By focusing on specific cell types or families of molecules, ICELLNET provides several representation modes to help the interpretation of the results. Experimentally validated with an *in vitro* system, ICELLNET enables the dissection of intercellular communication in complex systems.

## Methods

### Human Primary Cell Atlas dataset

The dataset contains 745 samples of over thirty human primary cell types in different biological conditions (rested or activated). Included in BioGPS platform, all the samples have been generated with the same Affymetrix technology (Human Genome U133 Plus 2.0 arrays). For this study, an already processed and normalized dataset has been downloaded and added to ICELLNET package.

#### CAFs RNA-seq data processing

The dataset contains 77 samples from Luminal (Lum) and Triple Negative Breast Cancers (TNBC) from 16 patients (10 Lum, 6 TNBC)^19^. The samples correspond either to tumor tissue or juxtatumoral tissue. Cells corresponding to CAF-S1 and CAF-S4 have been isolated, collected, and sequenced. Average sequencing depth was 30 million for paired-end reads, with a read length of 100bp. Reads were mapped on the reference genome (hg19/GRCh37 from UCSC genome release) using Tophat_2.0.6 algorithm. Duplicates were removed and gene expression quantification was performed using HTSeq-Count and featuresCount. Only genes with five reads in at least 25% of all samples were kept for further analyses. Normalization was done using the method implemented in DESeq2 R package. In this study, only 6 samples of CAF-S1 and 3 samples of CAF-S4 from TNBC were considered in the analyses.

#### Purification of Peripheral blood mononuclear cells (PBMCs) from adult blood

Fresh blood samples were collected from healthy donors and obtained from Hôpital Crozatier Établissement Français du Sang (EFS), Paris, France, in conformity with Institut Curie ethical guidelines. In agreement with EFS rules, all informed consent and consent to publish were obtained. PBMCs were isolated by centrifugation on a Ficoll gradient (Ficoll-Paque PLUS, GE Healthcare Life Sciences).

#### Monocyte-derived dendritic cells generation and activation

Monocytes were selected from PBMCs using antibody-coated magnetic beads and magnetic columns according to manufacturer’s instructions (CD14 MicroBeads, MiltenyiBiotec). To generate immature DCs, CD14+ cells were cultured for 5 days with IL-4 (50 ng/mL) and GM-CSF (10 ng/mL) in RPMI 1640 Medium, GlutaMAX (Life Technologies) with 10% FCS. Monocyte-derived DCs were pre-treated for one hour with mouse IgG1 (20 µg/mL, R&D Systems), mouse anti-IL-10R blocking antibody (10 µg/mL, R&D Systems) or mouse anti-TNFα Receptors 1 and 2 (10 µg/mL, R&D Systems) (see Figure 1-Figure Supplement 4B) and then cultured with medium or LPS (100 ng/mL, LPS-EB Ultrapure, activates TLR4 only, Invivogen) for 24 hours. DCs from donors which responded to (a) LPS and (b) IL-10R blocking antibody, as evaluated by maturation markers, were included in this study. The following cytokines were measured in culture supernatants by CBA (BD Bioscience): IL-6, IL-12p70 and OSM. IL-23 was measured using ELISA (eBioscience).

#### DC gene expression profiling

Monocyte-derived DCs were pre-treated with blocking Abs as described above for one hour and then cultured with medium or LPS (100 ng/mL, Invivogen) for an additional 4 or 8 hours. Total RNA was extracted using the RNeasy micro kit (Qiagen). Samples were then amplified and labelled according to the protocol recommended by Affymetrix for hybridization to Human Genome U133 Plus 2.0 arrays. If multiple probes corresponded to the same receptor, we selected the optimal probe based on the Jetset optimality condition^48^.

#### Curation of the ligand/receptor database

Surveying the literature for any potential interactions, we manually curated a ligand-receptor database using STRING (http://string-db.org/), Ingenuity (http://www.ingenuity.com/) and BioGRID (https://thebiogrid.org) online tools to verify protein-protein interactions, as well as Reactome and CellPhoneDB databases, already dedicated to ligand-receptor interactions. The interactions were classified into families of molecules based on the known biological function of the ligand and the receptor. The subfamilies of cytokines were defined based on molecular structures, as defined in the literature^3,14–16^. The database of ligand-receptor interactions is contained in the **supplementary table 1**.

#### Gene expression matrix scaling method

After selecting the genes corresponding to the ligands and/or receptors from the transcriptional profiles, each ligand/receptor gene expression is scaled among all the conditions ranging from 0 to 10. For each gene, the maximum value (10) is defined as the mean of expression of the 5% highest values of expression, and the minimum value (0) is defined as the mean expression of the 5% lowest values of expression. Outliers are rescaled at either 0 (if below minimum value) or 10 (if above maximum value).

#### Intercellular communication score computation

To score the intensity of a particular ligand-receptor interaction between a central cell and a given peripheral cell, we considered the product of the expression of the ligand in the central cell and of the cognate receptor in the peripheral cells. Formally, if 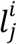 is the average expression level of ligand *i* by the central cell in the experimental condition *j*, and 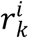 is the average expression of the corresponding receptor by cell type *k*, the intensity 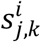 of the corresponding interaction was quantified by 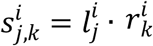. For interactions requiring multiple components of the ligand and/or of the receptor, we considered a geometric average of the receptor components. For example, if a given interaction corresponding to ligand *i* required two chains of the receptor, the score was computed as 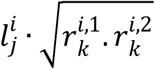, where 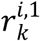 and 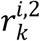 are the expression levels of the two receptor chains in cell type *k*. To assign a global score *S*_*j,k*_ to the communication between the central cell in the condition *j* and cell type *k*, a composite score was defined by summing up the intensity of all the possible ligand-receptor interactions, i.e., 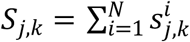, being the total number of interactions. Regarding the four DC experimental conditions (Medium (j=0), LPS (j=1), blocking TNF loop (j=2), blocking IL-10 loop), we normalized the global scores *S* _*j,k*_ to the Medium condition (*j*=0) across the four conditions. Thus, the final scores 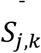 used to measure the communication intensity between DC in the condition *j* and the target cell *k* were computed using the following formula 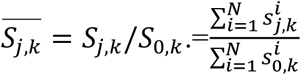. The score corresponding to each interaction and each target cell in the experimental condition of CAF subsets and the four DC experimental conditions are provided in **supplementary table 3A-C and 4A-E** respectively. The generation of the inward connectivity maps was done by reversing the role of the central cell and their cellular targets.

#### Global intercellular communication score scaling method

The intercellular communication scores are rescaled ranging from 1 to 10, considering all the scores computed for each biological condition between the central cell and all selected peripheral cell types. This step allows us to increase the differences between the scores and facilitate the network visualisation of the communication scores.

#### Statistical comparison of communication scores

To compare the communication scores obtained from the same central cell to different peripheral cells we compute several communication scores considering the average expression of ligands for the central cell and each replicate separately for the receptor expression of the peripheral cells. In this way, for one peripheral cell type, we obtain a distribution of n communication scores, n being the number of peripheral cells replicates for this particular cell type. Second, we can compare communication scores between two biological conditions. In this case, we compute several communication scores considering each replicates of the central cell separately, and the average gene expression for the peripheral cells. We obtain a distribution of n communication scores, n being the number of central cell replicates in one biological condition. For both cases, we then perform a Wilcoxon statistical test to compare the communication scores distributions. The p-values are adjusted with p.adjust() function from the R package « stats » (version 3.6.1) using the Benjamini & Hochberg^49^ method in R. This returns the p-value matrix of statistical tests, that can be visualized in a heatmap representation with the pvalue.plot() function from « icellnet » R package.

#### Statistical analysis of gene expression data

Expression data were normalized with Plier. Transcriptomics analysis was performed in Matlab. For independent filtering, we used the function *geneverfilter*, which calculates the variance of each probe across the samples and identifies the ones with low variance. Probes with variance less than the 40^th^ percentile were filtered out. Differential analysis was performed using an ANOVA test (function *anova1*) at 4 hours and 8 hours. p-values were adjusted for multiple testing using the Benjamini-Hochberg correction using the function *mafdr*. Adjusted p-values <5% were considered significant (see **Suppl. Table 4F**).

#### Purification of naive CD4+ T lymphocytes

CD4^+^T lymphocytes were purified from PBMCs by immunomagnetic depletion with the human CD4^+^T cell Isolation KitII (MiltenyiBiotec), followed by staining with allophyco-cyanin-anti CD4 (VIT4; MiltenyiBiotec), phycoerythrin-anti-CD45RA (BD), fluorescein-isothiocyanate-anti-CD45RO (BD Bioscience) and phycoerythrin-7-anti-CD25 (BD bioscience). Naive CD4^+^T cells sorting of CD4^+^CD45RA^+^CD45RO^-^CD25^-^ and memory CD4^+^ T cells sorted as CD4^+^CD45RA^-^ CD45RO^+^CD25^-^ had a purity of over 99% with a FACSAria (BD Bioscience).

#### DC-T cells Coculture assays

To analyze T cell polarization, 24 hours activated DC and T cells were incubated in 96 well plates at a DC/T ratio 1:5 in Xvivo15 medium (Lonza). After 6 days, T cells were resuspended in fresh Xvivo15 medium at a concentration of 1 million cells per mL and restimulated with anti-CD3/CD28 beads (life Technologies) at a ratio bead/cell 1:1. Supernatants of T cells were collected after 24 hours of restimulation. The following cytokines were measured in naive culture supernatants by Cytometric Bead Array (CBA) (BD Bioscience) according to the manufacturer’s instructions: IL-2, IL-3, IL-4, IL-9, IL-10, IL-17A, IL-17F and IFN-γ. Additional cytokines were measured in memory T cells supernatant: IL-5, IL-13, TNF and GM-CSF. Cytokine-producing cells were analyzed by intracellular staining after addition of brefeldinA (10ug/mL) during the last 3 hours of the 5 hours restimulation in PMA and ionomycine (100ng/mL and 500ng/ml respectively). Cells were stained for 30 minutes with the yellow live dead kit (Invitrogen). Finally, cells were fixed and permeabilized using the Staining Buffer Set (eBioscience) and stained with anti-IL9, anti-IFNg, and anti-IL17A (ebioscience), and analyzed by flow cytometry (BD Fortessa).

#### Measurement of surface molecules expression by plasmacytoid dendritic cells

In order to enrich plasmacytoid dendritic cells (pDCs), cells expressing CD3, CD9, CD14, CD16, CD19, CD34, CD56, CD66b and glycophorin A were depleted from PBMCs using magnetic sorting (Human Pan-DC Pre-Enrichment Kit, StemCell Technologies). pDCs were then sorted on a FACS Vantage instrument (BD Biosciences). pDCs were cultured for 24 hours at 37°C and 5% CO_2_ with medium RPMI 1640 Medium, GlutaMAX (Life Technologies) with 10% FCS, GM-CSF (10 ng/mL) used as a positive control or DC supernatants. Cells were stained for 15 minutes at 4°C using a FITC-anti-CD86 (BD), an APC-anti-ICOSL (R&D Systems) and Alexa-Fluor-700-anti-HLA-DR (Biolegend) or with the corresponding isotypes. Cells were analyzed on an LSR II instrument (BD Biosciences).

#### Measurement of adhesion molecules expression at the Neutrophil surface

Whole-blood samples were obtained from healthy donors from Hôpital Crozatier Établissement Français du Sang (EFS), Paris, France, in conformity with Institut Curie ethical guidelines. Blood samples were stimulated for an hour at 37°C with medium, LPS (100 ng/mL) used as a positive control or DC supernatants. Cells were stained at 4°C for 15 minutes with an APC-anti-Human-CD62L (clone DREG-56, BD Pharmingen), a BV650-anti-Human-CD11b (BioLegend) and a PE-anti-Human-CD15 (MiltenyiBiotec) or with the corresponding isotypes. Erythrocytes were lysed with 1X BD Pharm Lyse Solution (BD Pharmingen). White cells were resuspended in PBS supplemented with 1% human serum and 2 mM EDTA and analyzed on an LSR Fortessa instrument (BD Biosciences).

#### Real-time quantitative RT-PCR

The keratinocyte cell line HaCaT was cultured in DMEM (Gibco) supplemented with 10% FBS and 1% penicillin/streptomycin. Cells were cultured with medium, LPS (100 ng/ml), or with DC supernatant diluted 1:10 for 4 hours. Total RNA was extracted by RNeasy Mini kit (Qiagen). RNA was then transcribed to cDNA using Superscript II reverse transcriptase based on the manufacture’s protocol (Invitrogen). The Taqman method was used for real-time PCR with primers from Life technologies. The expression of mRNA was normalized to the geometrical mean of 3 house-keeping genes: β-actin, GAPDH and RPL34. All HaCaT cells were negative for Mycoplasma contamination, standardized and regular tests were performed by PCR for mycoplasma detection.

#### Statistical analysis of DC-T cell protein data

All analyses were generated with R 3.1. For principal component analysis (PCA) of the T cell secretion profile, a data matrix was formed whose rows corresponded to conditions and columns to the different cytokines (each column was scaled using *zscore*). PCA was done using the function *princomp*. Where appropriate, a paired student t-test was performed. Significant differences were considered with p<0.05. The correlation heatmap based on Spearman was generated on the logged data. Correlations with p values<0.05 were considered as significant.

#### Calculation of the activation score of target cells

To compute a global activation score of keratinocytes, neutrophils and pDC, each activation marker output was first normalized in the range 0-1, 0 being to the untreated condition and 1 being to the maximum value observed in all the conditions. An average of the normalized outputs corresponding to the same cell type was then considered. All of the measured factors, with the exception of CD62L in neutrophils, were positively correlated with cell activation. In order to make CD62L consistent with the other factors, we considered the reciprocal of its value. The numerical results are in the **Suppl. Table 5**.

## Data availability

The gene expression profiles generated for this publication have been deposited in NCBI’s Gene Expression Omnibus and are accessible through GEO Series accession number GSE89342 (http://www.ncbi.nlm.nih.gov/geo/query/acc.cgi?acc=GSE89342).

The CAFs dataset has been published by Costa et al. 2018^19^, and is accessible through the accession number EGAS00001002508.

## Code availability

ICELLNET package is available at https://github.com/soumelis-lab/ICELLNET

## Acknowledgements

We wish to thank Lilith Faucheux, Melissa Saichi, Fanny Coffin, Philippe Hupé, Franck Perez, Sebastian Amigorena, Yong-Jun Liu and Fivos Soumelis for insightful comments and discussions, the Institut Curie Flow Cytometry facility (Z. Maciorowsky), the Institut Curie Affymetrix facility (D. Gentien). This work was supported by funding from Institut National de la Santé et la Recherche Médicale (Inserm), the Institut Curie, Agence Nationale pour la Recherche (ANR), Fondation pour la Recherche Médicale (FRM), the European Research Council (ERC starting grant 281987), ANR-10-IDEX-0001-02 PSL, ANR-11-LABX-0043, CIC IGR-Curie 1428 for V.S. EMBO, Institut Curie post-doctoral fellowships to ICL. ANRS and ARC fellowships to MG and La ligue contre le cancer doctoral fellowship to LMR.

## Author contributions

F.N. and L.M.-R. designed and performed bioinformatic analyses and wrote the manuscript; I.C-L. designed and performed experiments and analyzed results; A.C performed bioinformatic analyses; M.G. and C.T. performed experiments and revised the manuscript; Y.K. and F.M.-G. provided strategic advices and revised the manuscript; and V.S. designed experiments, supervised the research and wrote the manuscript.

