## Supplementary figures for "ICELLNET: a transcriptome-based framework to dissect intercellular communication"

### Supplementary figures legends

**Supplementary Figure S1: Comparison of cytokine-mediated intercellular communication between Triple-Negative breast cancer infiltrating CAF subsets CAF-S1 and CAF-S4. (A)** Statistical score comparison of the outward cytokine-mediated communication scores computed from CAF-S1 (n=6) to the different peripheral cells (left), and the outgoing communication scores from CAF-S4 (n=3) to the different peripheral cell types (right). (**B)** Individual outward communication score of specific ligand-receptor interactions that contribute to communication scores from CAF-subsets to breast fibroblasts with a score superior or equal to 20.

**Supplementary Figure S2: Dissecting inward communication of Triple Negative CAF-subsets from putative cell types present in the tumor microenvironment (A)** Connectivity maps describing inward cytokine-mediatedcommunication from primary cells to CAF-S1 (n=6) and CAF-S4 (n=3) subsets. The CAF subsets are considered as central cells and colored in grey. Primary cells are considered as peripheral cells and are colored depending on the cell compartment (green: stroma, orange: innate, blue: adaptive, pink: epithelium). The width of the edges corresponds to a global score combining the intensity of all the individual ligand-receptor interactions. A scale ranging from 1 to 10, corresponding to minimum and maximum communication scores, is shown in the legend. **(B)** Barplot of communication score with contribution restricted to cytokines subfamilies between CAF subsets and a selection of peripheral cells.

**Supplementary Figure S3: IL-10R blocking activates a cell-to-cell communication module in LPS-stimulated DCs. (A)** Cell viability of DC cultured 24 hours in the indicated blocking conditions (Medium, LPS activation, LPS and TNFR or IL-10R blocking antibodies) was assessed by DAPI staining (n=6). Histograms represent the mean ± SEM of DAPI negative cell percentage. **(B)** Barplot of communication score with contribution by families of communication molecules between in vitro activated DCs in each biological condition and a selection of peripheral cells.

**Supplementary Figure S4: Observed effect on communication partner-cells is not due to the**

**presence of residual $\alpha$IL-10R antibody or a potent LPS dose.** (**A**) CD4 Naive T cells were pre-treated with blocking antibody for IL-10 receptor or a non-specific one and then put in culture with DC as indicated for 6 days. After restimulation with anti-CD3/anti-CD28 for 24 hours, supernatants were analyzed for the presence of IL-17F. Histogram represent means ± SEM (n= 4 donors). **(B)** HaCat cells were pre-treated with blocking antibody for IL-10 receptor or a non-specific one and then put in culture with DCs supernatant (diluted 1:10) as indicated for 4 hours. RNA was then extracted from cells and the expression of TNF and IL-1b was assayed using qRT-PCR (n=4 replicates). **(C)** pDCs were pre-treated with blocking antibody for IL-10 receptor or a non-specific one and then put in culture with DCs supernatant (diluted 1:10) as indicated for 24h. Expression of maturation markers CD86 and ICOSL analyzed by flow cytometry. (n= 6 donors). **(D)** Neutrophils cultured for 1h with 1ng/ml LPS were not significantly activated compared to medium as assessed by surface expression of CD11b and CD62L by flow cytometry. (n=3).

**Supplementary table legends:**

***Supplementary Table 1***: Manually curated database of ICELLNET ligand-receptor interactions. The database is structured as following: column 1-5, gene symbol for ligand/receptor subunits, column 6, possible aliases for ligands and receptors, column 7-9, classification into families of molecules, column 10, PubMed ID corresponding to the interaction.

***Supplementary Table 2:*** Description of the BioGPS dataset integrated in ICELLNET package. It contains a total 745 transcriptomic profiles among 31 cells types generated with the same Affymetrix technology.

***Supplementary Table 3***: **(A)** CAF subsets outward communication scores considering all the ligand-receptor database, including raw global communication score, global communication score after rescaling, communication score of each family of molecules, and statistical analysis. **(B)** CAF outward individual communication scores with selected peripheral cells. **(C)** CAF subsets outward cytokine-mediated communication scores, including raw global communication score, global communication score after rescaling, communication score of each subfamily of cytokines, and statistical analysis.

***Supplementary Table 4***: **(A)** Global outward communication score from the in vitro-activated DCs to selected peripheral cells, including raw global scores, and global scores after rescaling, steps. **(B-E)** List of individual outward communication scores from each DCs biological condition to selected peripheral cells. **(F)** Differential expression analysis was performed using an ANOVA test (Matlab function anova1) for these two time points. P-values were adjusted for multiple testing using the Benjamini-Hochberg correction using the Matlab function mafdr. Adjusted p-values <5% were considered significant.

***Supplementary Table 5:*** Expression of maturation markers CD86, HLA-DR and ICOSL or CD11b and CD62L analyzed by flow cytometry with surface staining on **(A)** pDCs (n=18), **(B)** keratinocytes (n=8), and **(C)** neutrophils (n=9) cultured with supernatant of the indicated DC.

Figure S1

A

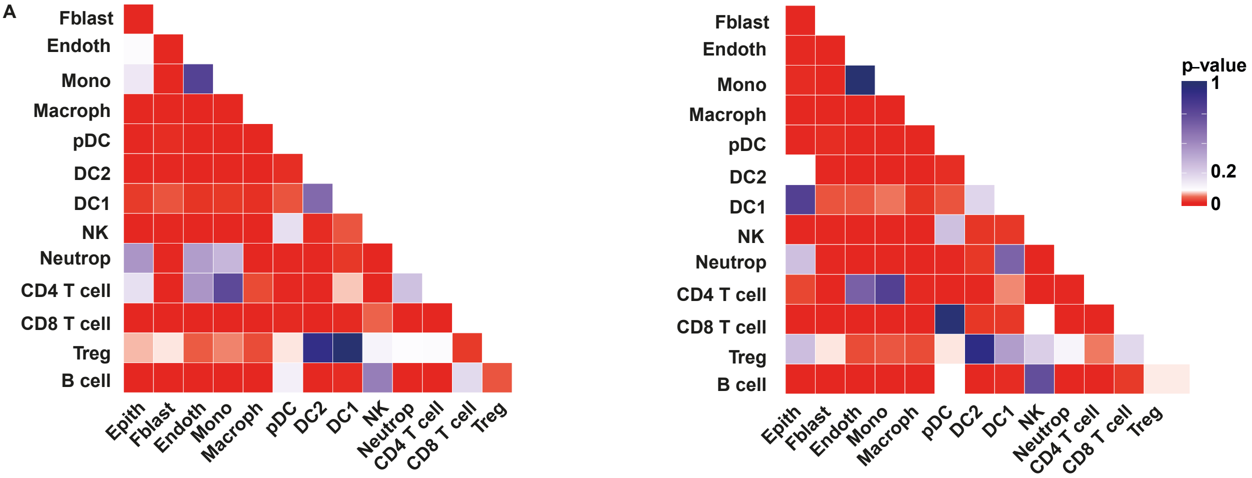
Fblast
Endoth
Mono
Macroph
pDC
DC2
DC1
NK
Neutrop
CD4 T cell
CD8 T cell
Treg
B cell
Epith
Fblast
Endoth
Mono
Macroph
pDC
DC2
DC1
NK
Neutrop
CD4 T cell
CD8 T cell
Treg
p-value
1
0.2
0

B

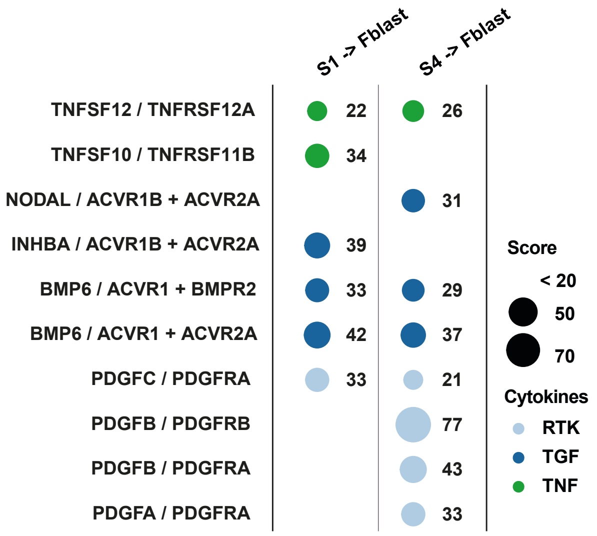
S1 -> Fblast
S4 -> Fblast
TNFSF12 / TNFRSF12A
TNFSF10 / TNFRSF11B
NODAL / ACVR1B + ACVR2A
INHBA / ACVR1B + ACVR2A
BMP6 / ACVR1 + BMPR2
BMP6 / ACVR1 + ACVR2A
PDGFC / PDGFRA
PDGFB / PDGFRB
PDGFB / PDGFRA
PDGFA / PDGFRA
22
26
34
31
39
33
29
42
37
33
21
77
43
33
Score
< 20
50
70
Cytokines
RTK
TGF
TNF

Figure S2

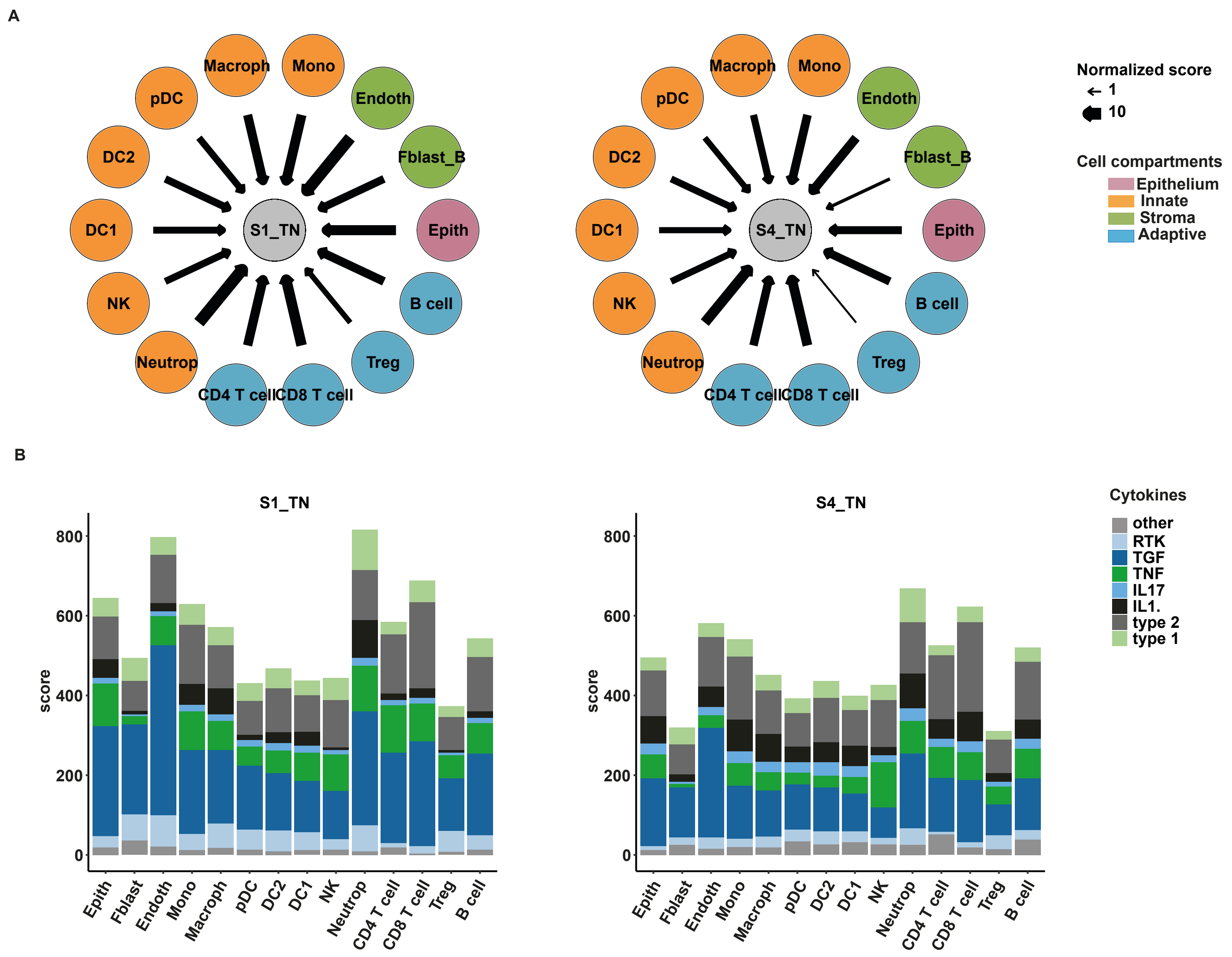
A
Macroph
Mono
pDC
Endoth
DC2
Fblast_B
DC1
S1_TN
Epith
NK
B cell
Neutrop
Treg
CD4 T cell
CD8 T cell
S4_TN
Normalized score
1
10
Cell compartments
Epithelium
Innate
Stroma
Adaptive
B
S1_TN
S4_TN
score
800
600
400
200
0
Epith
Fblast
Endoth
Mono
Macroph
pDC
DC2
DC1
NK
Neutrop
CD4 T cell
CD8 T cell
Treg
B cell
Cytokines
other
RTK
TGF
TNF
IL17
IL1.
type 2
type 1

Figure S3

A

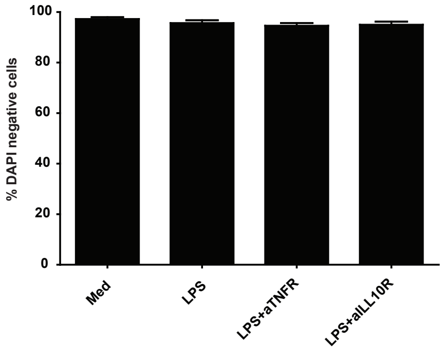
% DAPI negative cells
100
80
60
40
20
0
Med
LPS
LPS+aTNFR
LPS+aILL10R

B

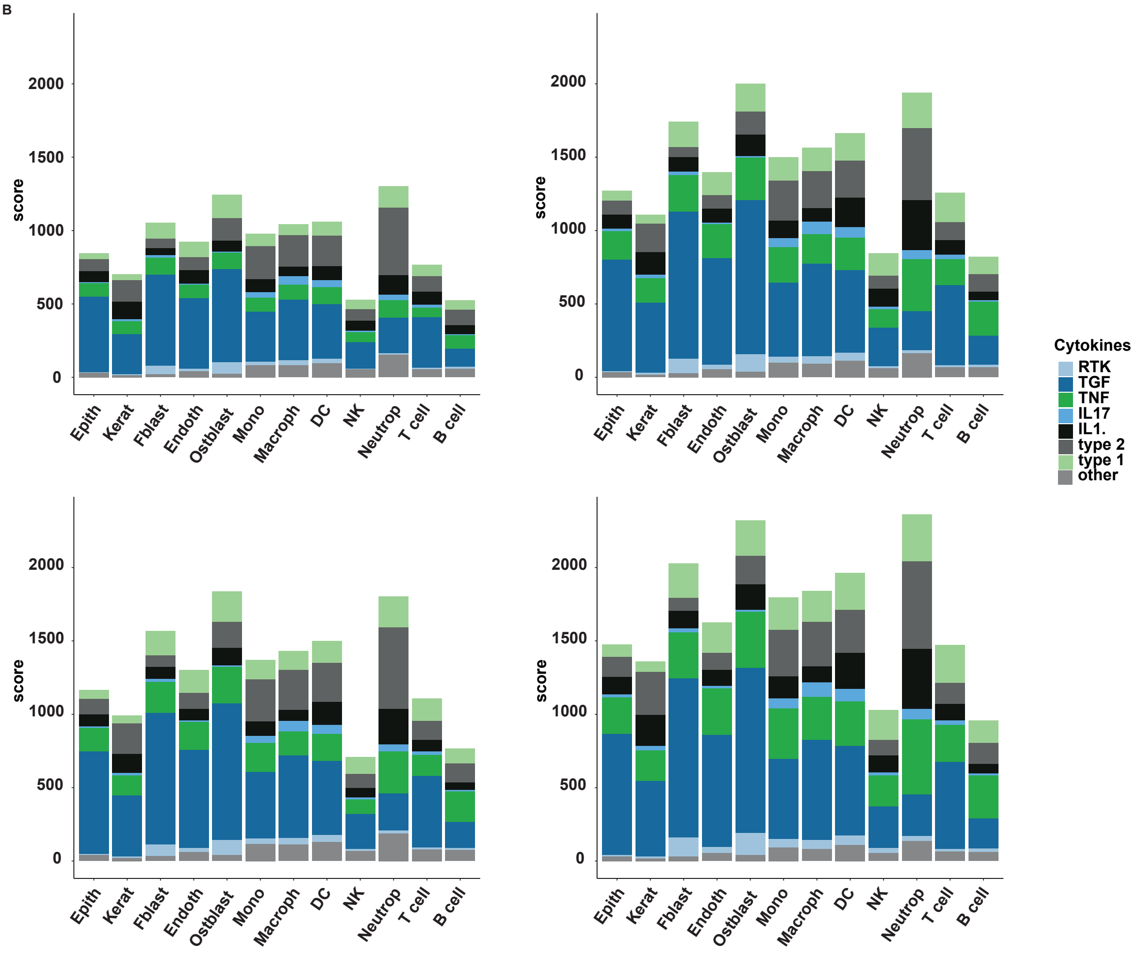
score
2000
1500
1000
500
0
Epith
Kerat
Fblast
Endoth
Ostblast
Mono
Macroph
DC
NK
Neutrop
T cell
B cell
Cytokines
RTK
TGF
TNF
IL17
IL1.
type 2
type 1
other

Figure S4

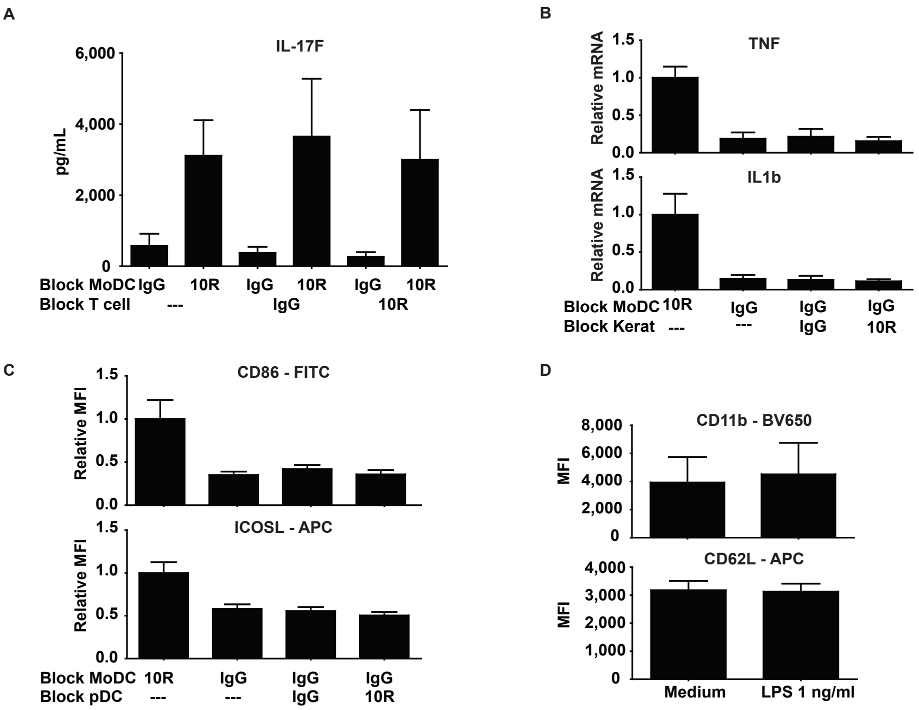
A
IL-17F
pg/mL
6,000
4,000
2,000
0
Block MoDC IgG 10R IgG 10R IgG 10R
Block T cell --- IgG 10R
B
TNF
Relative mRNA
1.5
1.0
0.5
0.0
IL1b
Relative mRNA
Block MoDC 10R IgG IgG IgG
Block Kerat --- --- IgG 10R
C
CD86 - FITC
Relative MFI
ICOSL - APC
Relative MFI
Block MoDC 10R IgG IgG IgG
Block pDC --- --- IgG 10R
D
CD11b - BV650
MFI
8,000
6,000
4,000
2,000
0
CD62L - APC
MFI
4,000
3,000
2,000
1,000
0
Medium
LPS 1 ng/ml
